# Chromosome-scale genome assembly of Japanese pear (*Pyrus pyrifolia*) variety ‘Nijisseiki’

**DOI:** 10.1101/2020.12.17.423204

**Authors:** Kenta Shirasawa, Akihiro Itai, Sachiko Isobe

## Abstract

**Aim:** The Japanese pear (*P. pyrifolia*) variety ‘Nijisseiki’ is valued for its superior flesh texture, which has led to its use as a breeding parent for most Japanese pear cultivars. However, in the absence of genomic resources for Japanese pear, the parents of the ‘Nijisseiki’ cultivar remain unknown, as does the genetic basis of its favorable texture. The genomes of pear and related species are complex due to ancestral whole genome duplication and high heterozygosity, and long-sequencing technology was used to address this.

**Methods and Results:** De novo assembly of long sequence reads covered 136× of the Japanese pear genome and generated 503.9 Mb contigs consisting of 114 sequences with an N50 value of 7.6□Mb. Contigs were assigned to Japanese pear genetic maps to establish 17 chromosome-scale sequences. In total, 44,876 protein-encoding genes were predicted, 84.3% of which were supported by predicted genes and transcriptome data from Japanese pear relatives. As expected, evidence of whole genome duplication was observed, consistent with related species.

**Conclusion and Perspective:** This is the first genome sequence analysis reported for Japanese pear, and this resource will support breeding programs and provide new insights into the physiology and evolutionary history of Japanese pear.

## 1. Introduction

Pear (*Pyrus* spp.) is a genus of the Malinae subtribe of the Rosaceae that includes European pear (*P. communis*), Chinese white pear (*P.* × *bretschneideri*), Japanese pear (*P. pyrifolia*), and apple (*Malus* × *domestica*). The Japanese pear variety ‘Nijisseiki’ (a Japanese term referring to 20^th^-century) was discovered in Matsudo (Chiba, Japan) in 1888^1^. Due to its favorable flesh texture, ‘Nijisseiki’ was the leading Japanese pear variety in Japan from the 1940s to the 1980s, and ‘Nijisseiki’ was widely used as a breeding parent for the development of Japanese pear cultivars in Japan^1^.

Genome information can enhance breeding programs^2^ by facilitating understanding of the genetic backgrounds of the breeding pedigrees^3^, and by enhancing physiology and evolutionary studies. In the Malinae, genome sequence data are publicly available for crop species such as apple^4^, European pear^5^, and Chinese white pear^6^, as well as for some pear wild relatives (*P. betuleafolia* and (*P. ussuriensis* × *communis*) × spp.) that are used for root stocks^7,8^. Genome analysis of the Malinae is complex due to ancestral whole genome duplication^9^ and high heterozygosity in their genomes resulting from allogamy and self-incompatibility. To simplify genomic analysis, doubled-haploid lines were developed to reduce the genome complexities in materials used for genome sequencing^5,6^. However, neither doubled-haploid lines nor genome sequence data are available for Japanese pear, despite the publication of transcriptome data^10^ and genetic maps11,12.

Long-read sequencing has several advantages over short-read technologies^13^. Long reads span repetitive sequence stretches in genomes, extending sequence contiguity and facilitating assembly. Furthermore, long reads allow haplotype phases of highly heterozygous genome sequences to be determined. In this study, long-read sequencing was used to produce a highly contiguous genome sequence assembly of the Japanese pear variety ‘Nijisseiki’. This Japanese pear genome will enhance our understanding of genetics, genomics, and breeding in Japanese pear.

## 2. Materials and methods

### 2.1. Plant materials and DNA extraction

A single tree of Japanese pear (*P. pyrifolia*), variety ‘Nijisseiki’, which is planted at the orchard of Kyoto Prefectural University (Kyoto, Japan), was used for genome sequencing analysis. Genome DNA was extracted from the young leaves by a modified sodium dodecyl sulfate method^14^.

### 2.2. Estimation of the genome size of Japanese pear

A short-read sequence library was prepared using a PCR-free Swift 2S Turbo Flexible DNA Library Kit (Swift Sciences, Ann Arbor, MI, USA) and converted into a DNA nanoball sequencing library with an MGI Easy Universal Library Conversion Kit (MGI Tech, Shenzhen, China). The library was sequenced on a DNBSEQ G400RS (MGI) instrument in paired-end, 101 bp mode. The obtained reads were used to estimate the genome size with Jellyfish after removing low-quality bases (<10 quality value) with PRINSEQ, adaptor sequences (AGATCGGAAGAGC) with fastx_clipper in FASTX-Toolkit, and reads from organelle genomes^15,16^ (GenBank accession numbers: AP012207 and KY563267) by read mapping with Bowtie2 on the reference sequences.

### 2.3. Chromosome-scale genome assembly

A long-read sequence library was constructed using an SMRTbell Express Template Prep Kit 2.0 and sequenced on SMRT cells (1M v3 LR) in a PacBio Sequel system (PacBio, Menlo Park, CA, USA). The obtained reads (≥15 kb) were assembled with Falcon, and the two haplotype sequences, primary contigs and haplotigs, of the diploid genome were resolved with Falcon-unzip. Potential sequence errors in the assembled sequences were corrected with ARROW using the PacBio reads. Haplotype duplications in the primary contigs were removed with Purge_Dups. Sequences from the organelle genomes, which were identified by a sequence similarity search of reported plastid^16^ (AP012207) and mitochondrial^15^ (KY563267) genome sequences from Japanese pear using minimap2, were also deleted. The final assembly was designated as PPY_r1.0. The primary contigs of PPY_r1.0 were assigned to genetic maps of Japanese pear^11,12^ with ALLMAPS, in which marker sequences were aligned on the contigs with BLAST. The resultant chromosome-scale pseudomolecule sequences were named PPY_r1.0.pmol. The haplotig sequences were aligned onto PPY_r1.0.pmol with minimap2. Completeness evaluation of the assembly was performed with BUSCO (embryophyta odb9). The chromosome-scale pseudomolecule sequences (PPY_r1.0.pmol) were compared with those of apple (GDDH13, v1.1)^4^, European pear (Bartlett DH, v2.0)^5^, and Chinese white pear (Dangshan Suli, v1.1)^6^ with D-GENIES. Whole genome duplications were detected with MCScanX, with threshold values of ≥85% sequence identity and ≤1e-100 E-values, and visualized with VGSC.

### 2.4. Gene prediction and repetitive sequence analysis

Protein-encoding genes were predicted by a MAKER pipeline using two training sets and a preset of SNAP for Arabidopsis. Peptide sequences of apple (GDDH13, v1.1)^4^, European pear (Bartlett DH, v2.0)^5^, and Chinese white pear (Dangshan Suli, v1.1)^6^ registered in the Genome Database for Rosaceae^17^, as well as transcriptome data for Japanese pear (Ppyrifolia_protein_v1.0)^10^, were also employed in the prediction. Short gene sequences of <300 bases, as well as genes predicted with an annotation edit distance of <0.5, were removed to facilitate selection of high-confidence genes. The genome positions of the high-confidence genes were compared with those of the peptide sequences from apple, European pear, Chinese white pear, and Japanese pear, and were aligned to PPY_r1.0.pmol in the MAKER pipeline. Functional annotation of the predicted genes was performed with Hayai-Annotation Plants.

Repetitive sequences in the PPY_r1.0.pmol assembly were identified with RepeatMasker, using repeat sequences registered in Repbase and a de novo repeat library built with RepeatModeler. The repeat elements were classified into nine types: short interspersed nuclear elements (SINEs), long interspersed nuclear elements (LINEs), long terminal repeat (LTR) elements, DNA elements, Small RNA, satellites, simple repeats, low complexity repeats, and unclassified, in accordance with RepeatMasker.

The software tools used for the data analyses are shown in Supplementary Table S1.

## 3. Results and data description

### 3.1. Assembly of the genome of a Japanese pear, ‘Nijisseiki’

In total, 76.9 Gb short-read data were used for estimation of the genome size of Japanese pear. *K*-mer distribution analysis indicated that the ‘Nijisseiki’ genome was highly heterozygous, and that the estimated haploid genome size was 537.1_Mb (Figure 1).

**Figure 1.**
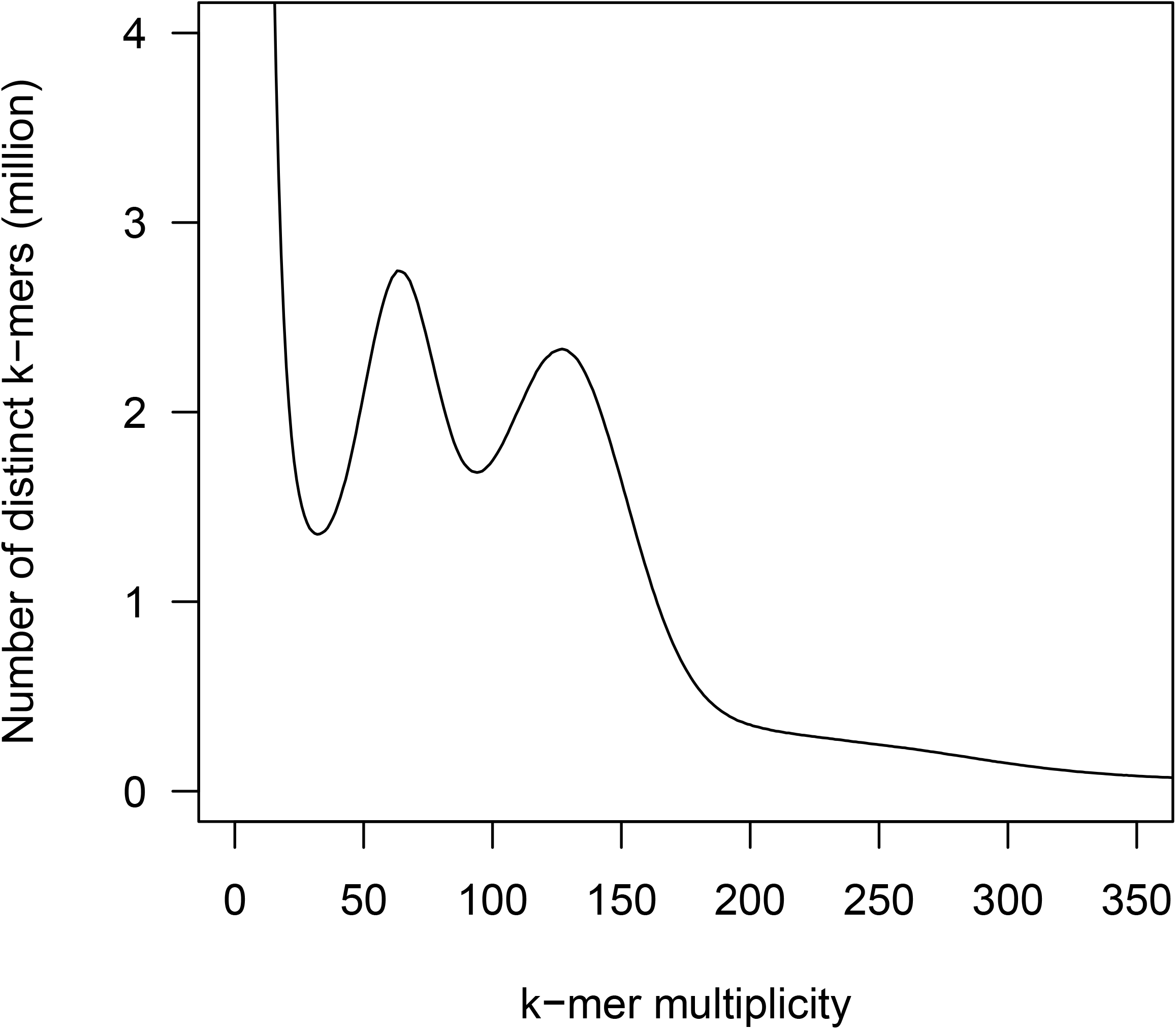
Genome size estimation for Japanese pear ‘Nijisseiki’ with the distribution of the number of distinct k-mers (k_=_17) with the given multiplicity values.

Long-read sequencing analysis with four SMRT cells generated 85.8 Gb data (4.5 million reads with an N50 of 25.5 kb). Of these, reads ≥15 kb in length (73.3 Gb, 136.4× genome coverage) were selected and assembled into 750.0 Mb contigs (495 sequences with an N50 of 5.9 Mb). The contig sequences were resolved into primary contigs and haplotigs, and polished to correct potential errors. The sizes of the resultant sequences were 745.8 Mb for the primary contigs and 113.3 Mb for the haplotigs. As this primary contig size was longer than the estimated haploid size, we doubted the existence of duplicate sequences. Hence, 257 haplotype duplications in the primary contigs (241.6 Mb) identified by a read depth distribution pattern of Purge_Dups (Supplementary Figure S1) were excluded. In addition, 10 sequences (283.5 kb), which showed significant similarity with previously reported organelle genomes, were removed. The final assembly, named PPY_r1.0, consisted of 503.9 Mb primary contigs (including 114 sequences with an N50 length of 7.6 Mb) and 353.4 Mb haplotigs (including 822 sequences with an N50 length of 1.6 Mb) (Table 1). A BUSCO analysis of the primary contigs indicated that 58.3% and 39.7% of sequences were single-copy and duplicated complete BUSCOs, respectively (Table 2), supporting the genome-wide duplication state in the Japanese pear genome.

**Table 1.** Statistics of the primary contig sequences of Japanese pear ‘Nijisseiki’

|  | PPY_r1.0<br>(primary contigs) | PPY_r1.0. haplotigs<br>(haplotig contigs) |
| --- | --- | --- |
| Total contig size (bp) | 503,888,133 | 353,451,501 |
| Number of contigs | 114 | 822 |
| Contig N50 length (bp) | 7,676,629 | 1,608,651 |
| Longest contig size (bp) | 28,225,311 | 7,932,431 |
| Gap (%) | 0 | 0 |
| #Predicted genes | 44,876 | Not available |

**Table 2.**
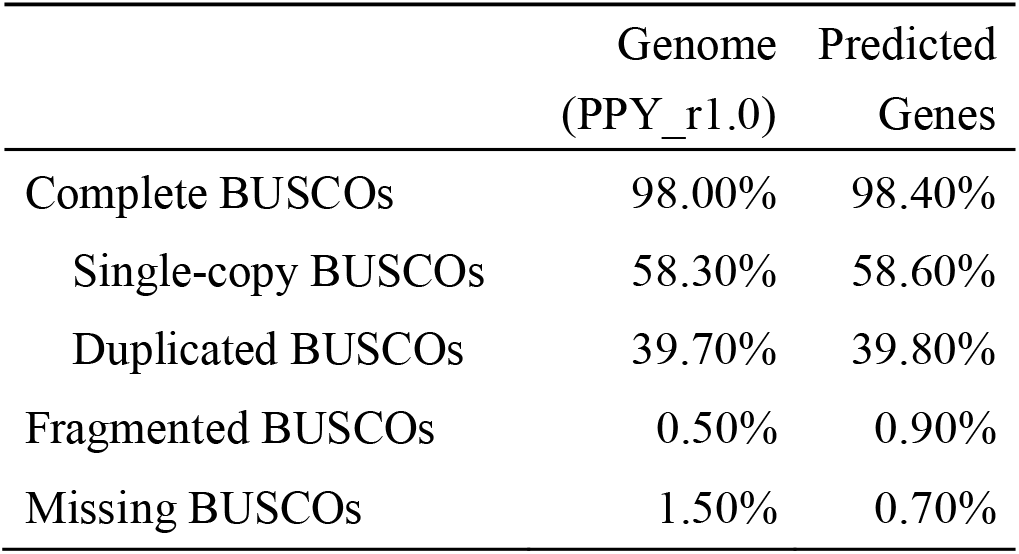
Completeness evaluation of genome assembly and predicted genes

|  | Genome<br>(PPY_r1.0) | Predicted<br>Genes |
| --- | --- | --- |
| Complete BUSCOs | 98.00% | 98.40% |
| Single-copy BUSCOs | 58.30% | 58.60% |
| Duplicated BUSCOs | 39.70% | 39.80% |
| Fragmented BUSCOs | 0.50% | 0.90% |
| Missing BUSCOs | 1.50% | 0.70% |

### 3.2. Chromosome-scale assembly construction and comparative genomics

Two previously established genetic maps for Japanese pear^11,12^ were used to establish pseudomolecule sequences. Totals of 609 and 2,388 SNP marker sequences were collected from the genetic maps of Li et al.^11^ and Terakami et al.^12^, respectively, and used to identify 587 and 2,306 marker positions on the primary contigs, respectively. Thus, 89 contig sequences (481.4 Mb in total) were assigned to the 17 linkage groups with the marker positions as anchors (Table 3), while the remaining 25 contigs (22.4 Mb) were not included. The haplotig sequences were aligned on the pseudomolecule sequences and covered 332.5 Mb in length. In total, 1,719,266 allelic sequence variants were detected between the two haplotypes.

**Table 3.** Statistics of the Japanese pear ‘Nijisseiki’ pseudomolecule sequences, PPY_r1.0.pmol

| Chromosome | #Contigs | % | Contig size<br>(bp) | % | #Genes | % |
| --- | --- | --- | --- | --- | --- | --- |
| Chr01 | 3 | 2.6% | 23,195,497 | 4.6% | 2,121 | 4.7% |
| Chr02 | 7 | 6.1% | 31,777,058 | 6.3% | 2,828 | 6.3% |
| Chr03 | 6 | 5.3% | 31,637,635 | 6.3% | 2,765 | 6.2% |
| Chr04 | 5 | 4.4% | 19,501,643 | 3.9% | 1,625 | 3.6% |
| Chr05 | 3 | 2.6% | 35,298,035 | 7.0% | 3,320 | 7.4% |
| Chr06 | 6 | 5.3% | 22,071,348 | 4.4% | 2,016 | 4.5% |
| Chr07 | 8 | 7.0% | 36,191,613 | 7.2% | 3,009 | 6.7% |
| Chr08 | 6 | 5.3% | 26,500,184 | 5.3% | 2,508 | 5.6% |
| Chr09 | 3 | 2.6% | 24,999,441 | 5.0% | 2,399 | 5.3% |
| Chr10 | 2 | 1.8% | 21,149,732 | 4.2% | 2,159 | 4.8% |
| Chr11 | 7 | 6.1% | 35,209,706 | 7.0% | 2,788 | 6.2% |
| Chr12 | 5 | 4.4% | 24,643,494 | 4.9% | 2,227 | 5.0% |
| Chr13 | 6 | 5.3% | 29,570,600 | 5.9% | 2,662 | 5.9% |
| Chr14 | 8 | 7.0% | 22,573,349 | 4.5% | 2,119 | 4.7% |
| Chr15 | 4 | 3.5% | 39,634,418 | 7.9% | 3,697 | 8.2% |
| Chr16 | 5 | 4.4% | 25,480,041 | 5.1% | 2,187 | 4.9% |
| Chr17 | 5 | 4.4% | 31,990,191 | 6.3% | 2,635 | 5.9% |
| **Subtotal**<br>**(Chr01-Chr17)** | **89** | **78.1%** | **481,423,985** | **95.5%** | **43,065** | **96.0%** |
| Unassigned<br>contigs | 25 | 21.9% | 22,747,703 | 4.5% | 1,811 | 4.0% |
| **Total** | **114** | **100.0%** | **504,171,688** | **100.0%** | **44,876** | **100.0%** |

The structure of the pseudomolecule sequences was compared with those of apple, European pear, and Chinese white pear to reveal one-to-one synteny relationships at chromosome level between the species (Figure 2). Furthermore, possible homologous chromosome pairs derived from genome-wide duplication and intra-chromosomal duplications were detected in all chromosomes (Figure 3, Table 4).

**Table 4.** Chromosome pairs showing sequence and structure homology in the Japanese pear genome

| Chromosome | Homologous chromosome<sup>a</sup> |
| --- | --- |
| 1 | 7, 15 |
| 2 | **2**, 7, 15, 17 |
| 3 | **3**, 11 |
| 4 | 12 |
| 5 | **5**, 10 |
| 6 | 14, 16 |
| 7 | 1, 2, **7** |
| 8 | **8**, 15 |
| 9 | 17 |
| 10 | 5 |
| 11 | 3 |
| 12 | 4, **12**, 14 |
| 13 | **13**, 16 |
| 14 | 6, 12 |
| 15 | 1, 2, 8 |
| 16 | 6, 13 |
| 17 | 2, 9 |

<sup>a</sup>Bold indicates intra-chromosome duplications.

**Figure 2.**
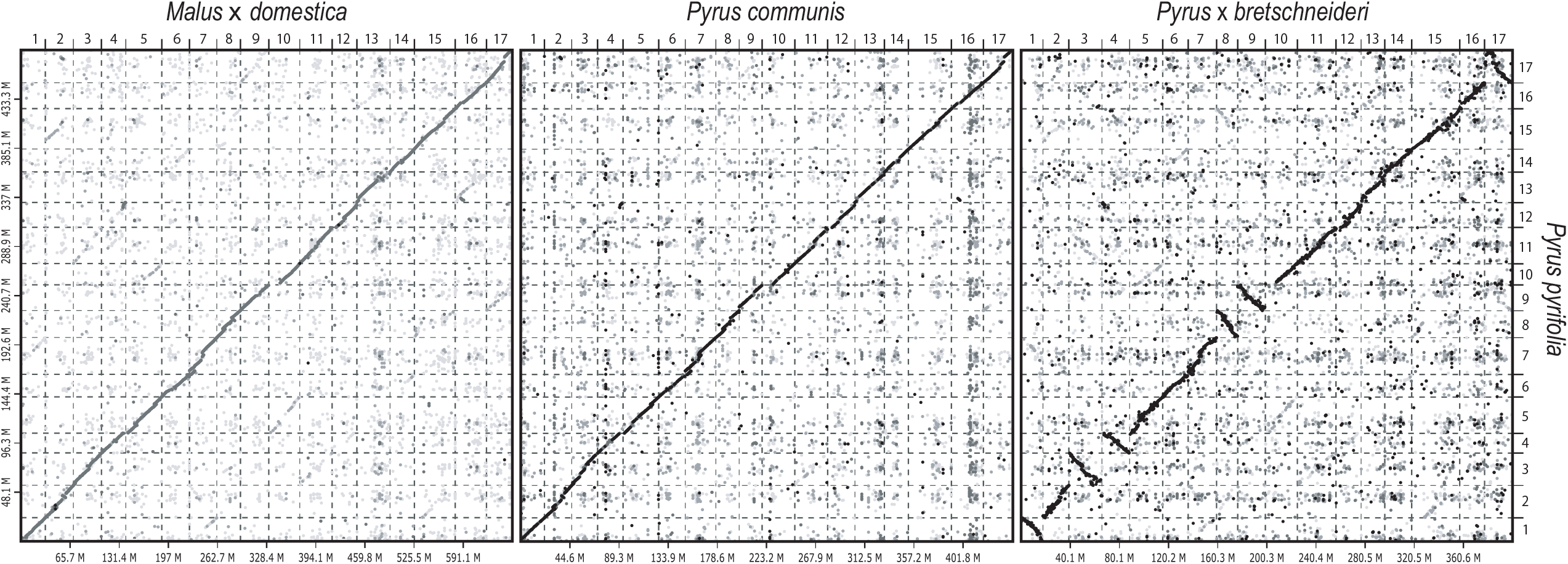
Comparative analysis of sequence and structure similarities of the Japanese pear genome. Genome sequence and structure similarities of Japanese pear (*P. pyrifolia*) with apple (*M.* × *domestica*), European pear (*P. communis*), and Chinese white pear (*P. bretschneideri*).

**Figure 3.**
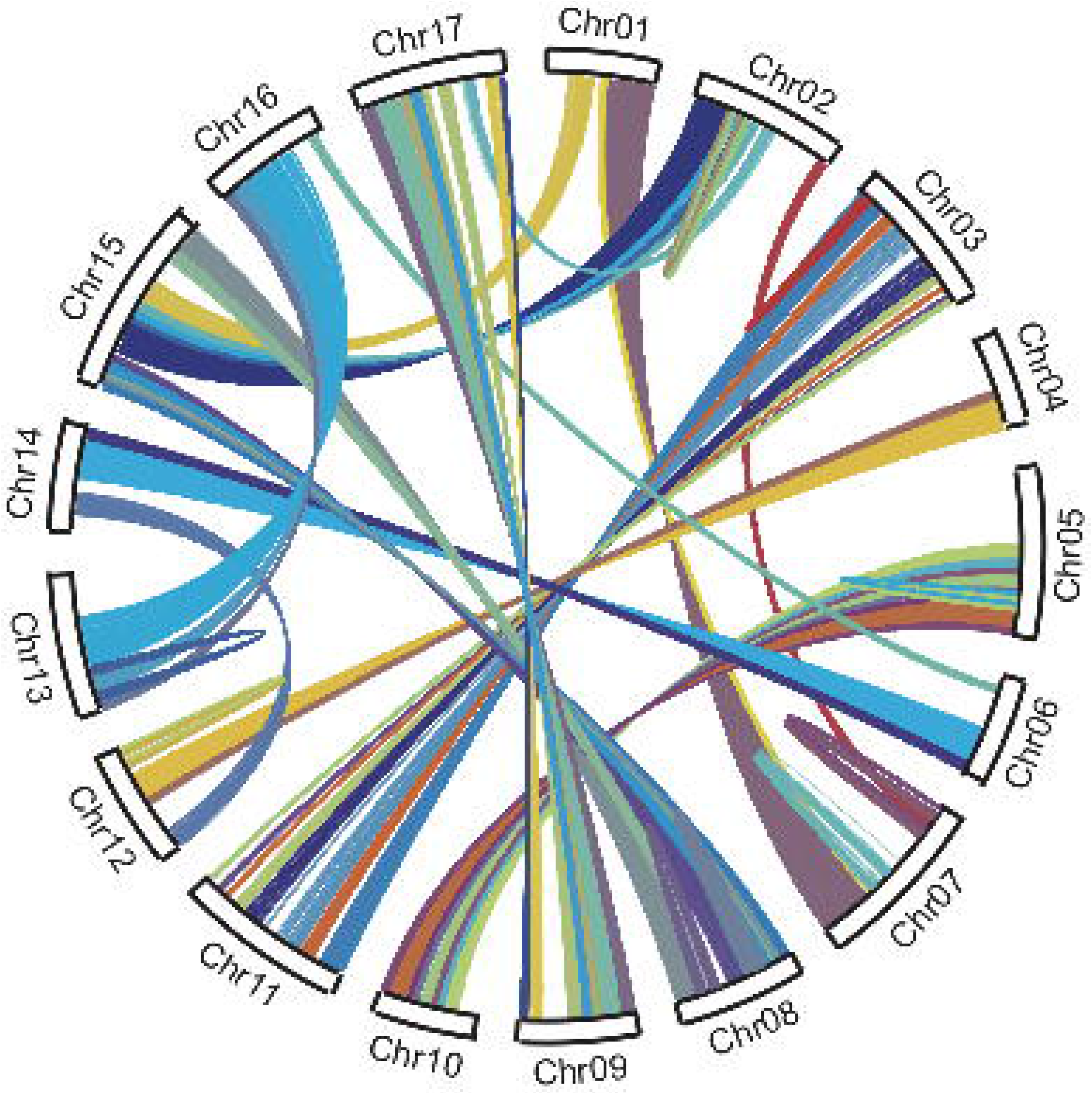
Whole genome duplication of the Japanese pear genome. Genome regions showing sequence similarities are connected with lines.

### 3.3. Gene prediction and repetitive sequence analysis

Initial predictions suggested 85,013 likely genes in the genome assembly. This gene number was resolved to 44,876 (Table 1) after removal of 29,607 low-confidence genes with an annotation edit distance of <0.5, 10,527 short genes of <300 bases in length, and three genes in redundant positions. A BUSCO analysis of the 44,876 remaining genes indicated that 98.4% of the sequences were complete BUSCOs (Table 2). The 44,876 sequences (56.0 Mb in total length) were therefore concluded to be high-confidence Japanese pear genes.

The genome positions of 37,835 (84.3%) of the 44,876 genes overlapped those of transcripts of Japanese pear and/or peptide sequences predicted from the genomes of apple, European pear, and Chinese white pear (Table 5). Furthermore, 6,610 (14.7%), 11,664 (26.0%), and 8,168 (18.2%) sequences were assigned to Gene Ontology slim terms in the biological process, cellular component, and molecular function categories, respectively, and 1,967 genes had enzyme commission numbers (Supplementary Table S2).

**Table 5.** Number of *P. pyrifolia* genes supported by other species of the Rosaceae

| Species supporting genes<sup>a</sup> | Number of genes | % |
| --- | --- | --- |
| Pb, Pc, Pp, and Md | 17,502 | 39.0% |
| Pb, Pc, and Pp | 735 | 1.6% |
| Pb, Pc, and Md | 4,603 | 10.3% |
| Pb, Pp, and Md | 1,737 | 3.9% |
| Pc, Pp, and Md | 1,832 | 4.1% |
| Pb and Pc | 699 | 1.6% |
| Pb and Pp | 437 | 1.0% |
| Pb and Md | 1,639 | 3.7% |
| Pc and Pp | 850 | 1.9% |
| Pc and Md | 1,350 | 3.0% |
| Pp and Md | 576 | 1.3% |
| Pb | 882 | 2.0% |
| Pc | 2,010 | 4.5% |
| Pp | 1,580 | 3.5% |
| Md | 1,403 | 3.1% |
| None | 7,041 | 15.7% |
| **Total** | **44,876** | **100.0%** |

<sup>a</sup>Pb: Chinese white pear (Dangshan Suli, v1.1)<sup>6</sup>; Pc: European pear (Bartlett DH, v2.0)<sup>5</sup>,
Pp: Japanese pear (Ppyrifolia\_protein\_v1.0)<sup>10</sup>; and Md: apple (GDDH13, v1.1)<sup>4</sup>.

Repetitive sequences occupied a total of 280.3_Mb (55.6%) of the PPY_r1.0.pmol (503.8_Mb) assembly. The repeats consisted of nine major types in varying proportions (Table 6). The dominant repeat types in the pseudomolecule sequences were LTR retroelements (172.9 Mb) followed by DNA transposons (59.4 Mb). Repeat sequences that were unavailable in public databases totaled 25.0 Mb.

**Table 6.**
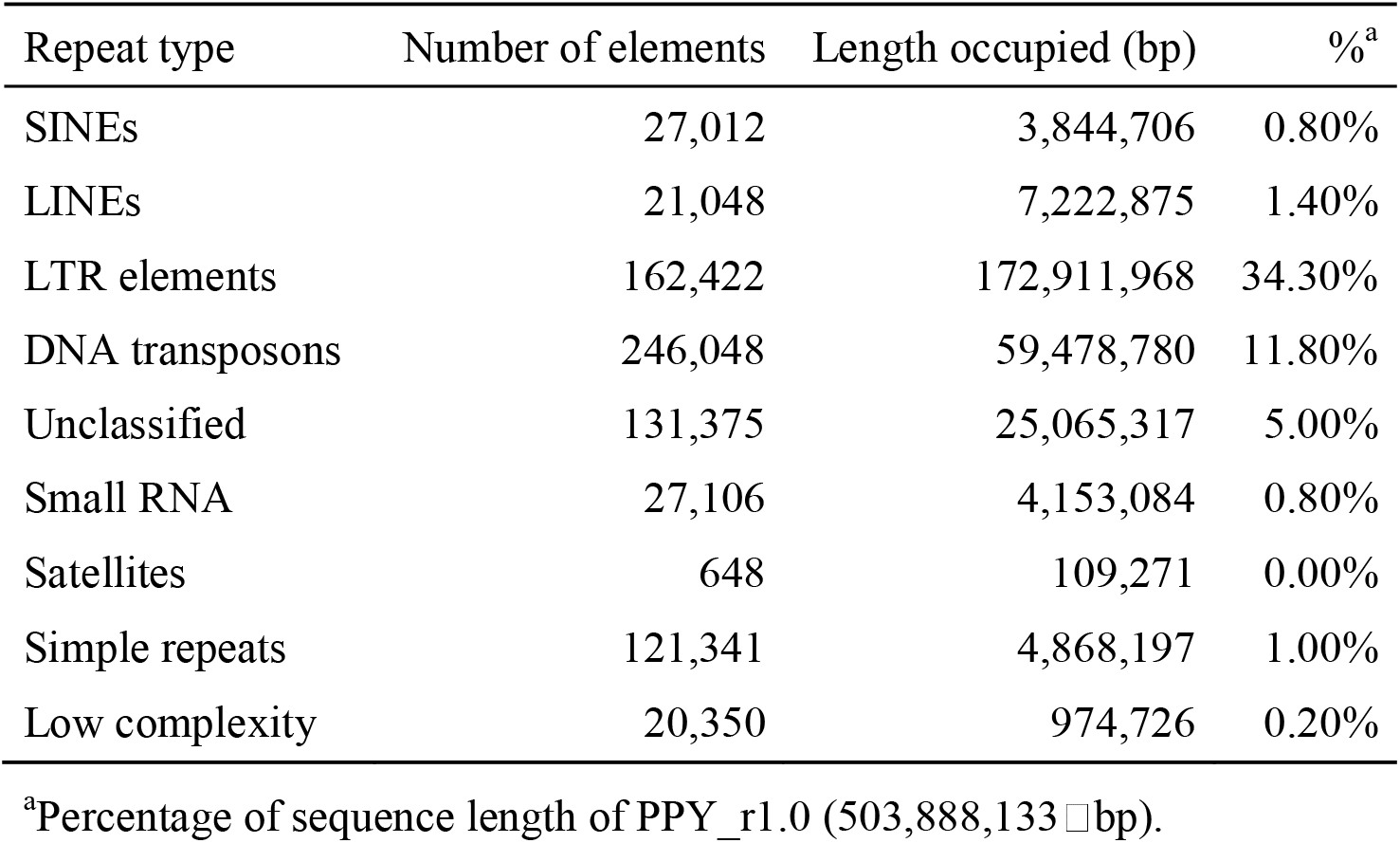
Repetitive sequences in the Japanese pear ‘Nijisseiki’ sequences, PPY_r1.0

| Repeat type | Number of elements | Length occupied (bp) | %<sup>a</sup> |
| --- | --- | --- | --- |
| SINEs | 27,012 | 3,844,706 | 0.80% |
| LINEs | 21,048 | 7,222,875 | 1.40% |
| LTR elements | 162,422 | 172,911,968 | 34.30% |
| DNA transposons | 246,048 | 59,478,780 | 11.80% |
| Unclassified | 131,375 | 25,065,317 | 5.00% |
| Small RNA | 27,106 | 4,153,084 | 0.80% |
| Satellites | 648 | 109,271 | 0.00% |
| Simple repeats | 121,341 | 4,868,197 | 1.00% |
| Low complexity | 20,350 | 974,726 | 0.20% |

<sup>a</sup>Percentage of sequence length of PPY\_r1.0 (503,888,133 bp).

## 4. Conclusion and future perspectives

Here, we present the first reported genome sequence analysis of Japanese pear, complementing the existing available genome sequences of other members of the *Pyrus* genus^5–8^. Long-sequencing technology allowed the assembly of 114 contig sequences (N50 of 7.6 Mb) spanning 503.9 Mb, 89.6% of the estimated size of the genome (Table 1). Subsequently, 481.4 Mb (95.5% of the assembled sequences) were successfully assigned to the pear chromosomes to establish pseudomolecule sequences, PPY_r1.0.pmol. This pseudomolecule assembly is the longest among the reported *Pyrus* spp.^5–8^, and PPY_r1.0.pmol can thus act as a new standard for *Pyrus* genomics. Genome structures and whole genome duplication events were conserved among pears and apples (Figure 2, Figure 3), despite their divergent genome sizes^2,4–8^. This Japanese pear genome assembly contributes to our understanding of the evolutionary history of pear species and the wider Malinae.

The ‘Nijisseiki’ variety of Japanese pear was used for the genome analysis. Although the parents of the ‘Nijisseiki’ variety are unknown, its favorable flesh texture^1^ led to its use as a popular breeding parent for Japanese pear cultivars. The genome data generated in this study will facilitate the discovery of the likely parents of the ‘Nijisseiki’ variety and will allow the ‘Nijisseiki’ haplotypes to be traced in progeny cultivars. As the superior flesh texture was unique to ‘Nijisseiki’ at the point of its discovery, and this texture was passed to progeny cultivars^1^, it should be possible to discover the genomic region responsible for the texture trait through pedigree analysis. This approach was used successfully to find associations between haplotypes and phenotypes for five traits in apple cultivars bred in Japan, for which the popular ‘Fuji’ variety was the founder^3^.

In summary, the Japanese pear genome was sequenced using long-read technology, and sequences were successfully assigned to the chromosomes. Genome structure was conserved among the members of the Malinae, and whole genome duplication was apparent in the Japanese pear genome. The Japanese pear genome will provide new insights into the physiology and evolutionary history of Rosaceae and will facilitate Japanese pear breeding programs.

## Acknowledgements

We thank Y. Kishida, C. Minami, H. Tsuruoka, and A. Watanabe (Kazusa DNA Research Institute) for their technical assistance.

## Data availability

Sequence reads are available from the Sequence Read Archive (DRA) of the DNA Data Bank of Japan (DDBJ) under accession number DRA011164. The DDBJ accession numbers of assembled sequences are BNSU01000001-BNSU01000114.

## Supporting information

**Supplementary Table S1.** Program tools used for genome assembly and gene prediction.

**Supplementary Table S2.** Functional annotations of predicted genes.

**Supplementary Figure S1.**Distribution of read depth on the primary genome assemblies.

A cutoff value at read depth of >117 to delete duplicated sequences is indicated by a vertical dotted line.

