## Supplementary Figure for "Chromosome-scale genome assembly of Japanese pear (*Pyrus pyrifolia*) variety ‘Nijisseiki’"

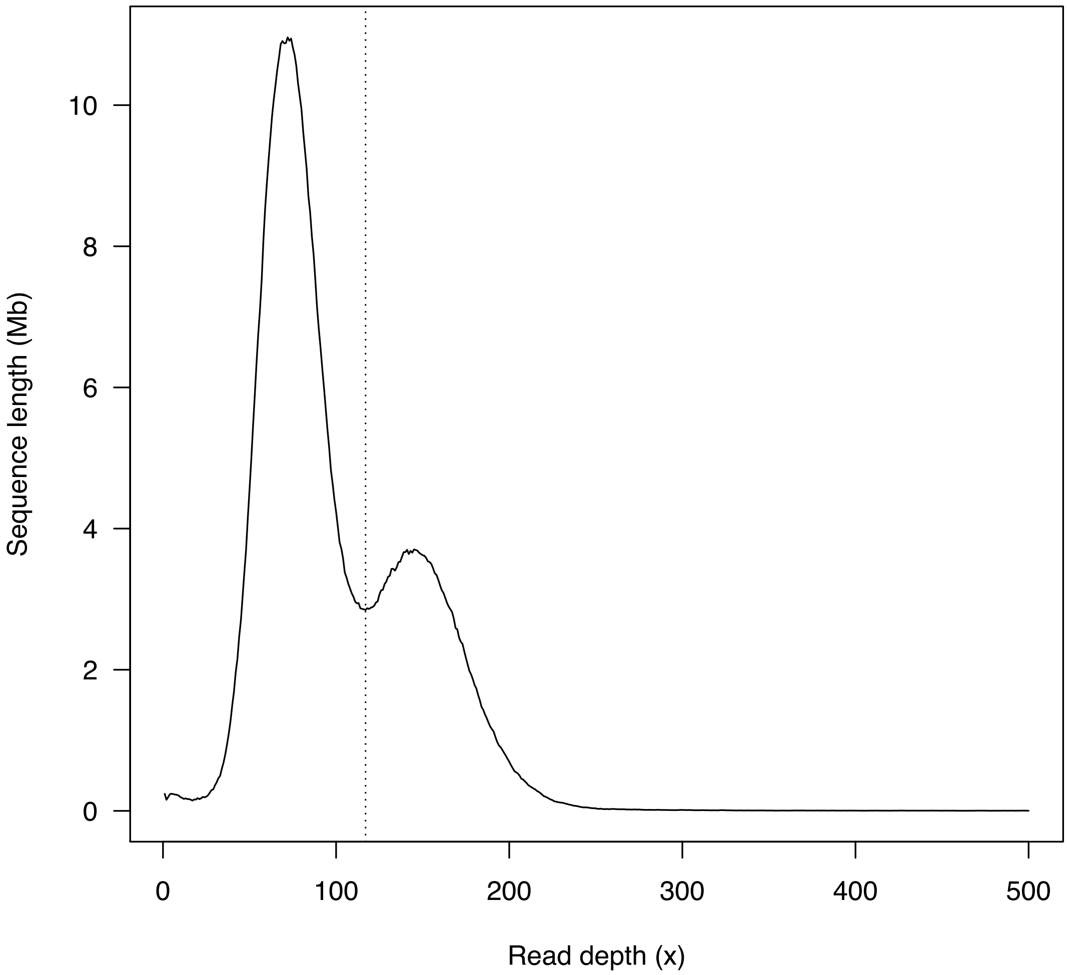


**Supplementary Figure S1** Distribution of read depth on the primary genome assemblies. A cutoff value at read depth of >117 to delete duplicated sequences are indicated by a dotted line.
